# CRISPR-based Live Imaging of Epigenetic Modification-Mediated Genome Reorganization

**DOI:** 10.1101/2020.02.18.954610

**Authors:** Ying Feng, Yao Wang, Chen Yang, Ardalan Naseri, Thoru Pederson, Jing Zheng, Xiao Xiao, Shaojie Zhang, Wei Xie, Hanhui Ma

**Author notes:** These authors contributed equally (Ying Feng, Yao Wang). Correspondence (W.X.), (H.M.).

## Abstract

Epigenetic modifications play an essential role in chromatin architecture and dynamics. The role of epigenetic modification in chromatin organization has been studied by Hi-C from population cells, but imaging techniques to study their correlation and regulation in single living cells are lacking. Here we develop a CRISPR-based EpiGo (Epigenetic perturbation induced Genome organization) system to track epigenetic modification-mediated relocation, interaction or reorganization of genomic regions in living cells. EpiGo-KRAB is sufficient to induce the relocation of genomic loci to HP1α condensates and trigger genomic interactions. EpiGo-KRAB also triggers the induction of H3K9me3 at large genomic regions, which decorate on the surface of HP1α condensates possibly driven by phase separation.

---

Human genome is organized in a hierarchy manner from kilobase to megabase scales such as nucleosome, loops, topologically associated domains (TADs) and A/B compartments ^1^. CTCF and cohesin play an important role in chromatin loop interaction ^2^. It has been proposed that loop extrusion drives TAD formation, while liquid-liquid phase separation mediates genome compartmentalization ^3^. For example, heterochromatin protein HP1α undergoes liquid-liquid demixing suggesting a role of phase separation in heterochromatin domain formation ^4, 5^. The co-segregated compartments often share similar chromatin states such as histone marks ^6^. Despite this widely observed correlations from Hi-C, whether the epigenetic modifications can indeed regulate genomic interactions and genome organization in single cells remains unclear. Direct visualization of chromatin structures in cell nucleus is still challenging. MERFISH, OligoSTORM or OCRA has been applied to trace DNA folding ^7–9^. These methods are applied to fixed cells. On the other hand, CRISPR-based imaging provides a versatile and powerful tool to track chromatin topology in live cells in real time ^10–12^.

Here we develop a CRISPR-based EpiGo system to investigate the effect of epigenetic modification on genome organization in living cells. We show EpiGo-KRAB induced H3K9me3 at different loci simultaneously, which mediated genomic interactions in single cells. We believe that this system should be applicable for exploring other epigenetic perturbations and genome organization.

The role of H3K9me3 in genomic interaction and compartmentalization were revealed by Hi-C in the population analysis, however direct visualization of genomic interaction-mediated by H3K9me3 in single living cells hasn’t been demonstrated. Here we developed a CRISPR-based system, namely EpiGo (Epigenetic perturbation induced Genome organization) (**Fig. 1A**) for live imaging of H3K9me3-mediated genomic interactions. We utilized dCas9-KRAB, which deposits loci-specific H3K9me3 modification ^13^, and fluorescent guide RNAs from the CRISPRainbow ^14^ or CRISPR-Sirius ^15^ system for genome visualization. We use HP1α as a reader for H3K9me3 ^4^ and HaloTag was knocked-in at the C-terminus of HP1α by the CRISPR-Cas9 system in U2OS cells, resulting U2OS-HP1α-HaloTag stable cell lines. The colocalization of H3K9me3 Loci and HP1α condensates were examined.

**Fig. 1.**
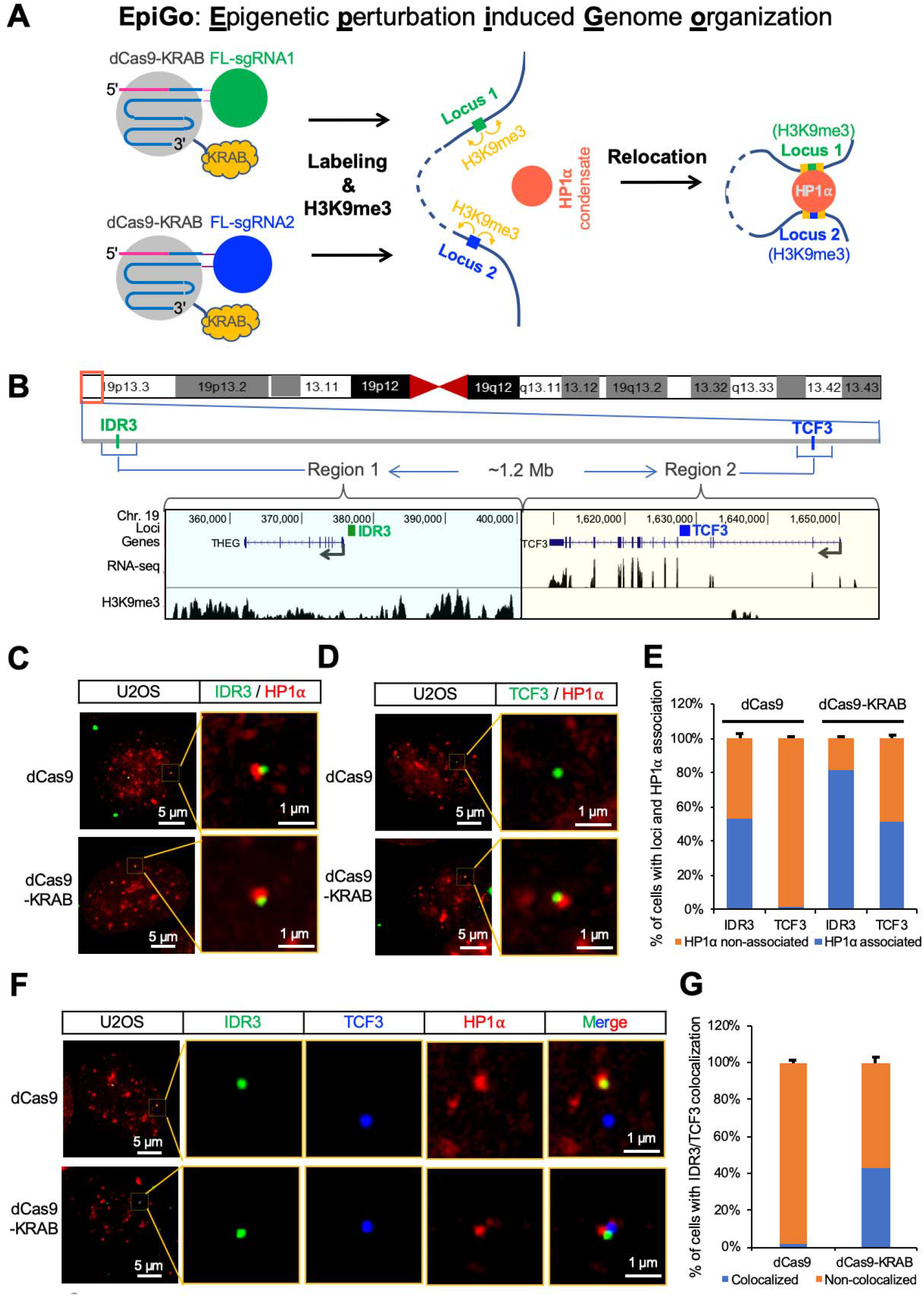
An CRISPR-based EpiGo system for live imaging of genomic interaction upon epigenetic modification. (A) A scheme of the EpiGo-KRAB system. EpiGo (Epigenetic perturbation induced Genome organization)-KRAB system consists of dCas9-KRAB, florescent sgRNAs such as FL-sgRNA1 and FL-sgRNA2 from either CRISPRainbow or CRISPR-Sirius targeting to genomic loci such as Locus 1 and Locus 2. dCas9-KRAB will induce H3K9me3 at each target sites or regions. Loci or regions with H3K9me3 will relocate to HP1α condensates, which will mediate genomic interactions. (B) RNA expression and H3K9me3 state of regions containing IDR3 or TCF3 loci in U2OS cells. RNA-seq and H3K9me3 ChIP-seq of two 50 kb regions were shown: Region 1 (chr19: 351,000-401,000) containing IDR3 and Region 2 (chr19: 1,605,500-1,655,500) containing TCF3. Region 1 and Region 2 are about 1.2 Mb apart on the p-arm of human chromosome 19. (C) Colocalization of IDR3 and HP1α in EpiGo-Control (dCas9) or EpiGo-KRAB (dCas9-KRAB) cells. IDR3 was imaged by CRISPR-Sirius in presence of PCP-GFP, IDR3-sgRNA-8XPP7 and dCas9 or dCas9-KRAB. U2OS-HP1α-HaloTag cells was used to examine the colocalization between IDR3 (green) and HP1α (red). (D) Colocalization of TCF3 and HP1α in EpiGo-Control (dCas9) and EpiGo-KRAB (dCas9-KRAB) cells. TCF3 was labeled and imaged with the same conditions as (C) except that IDR3-sgRNA-8XPP7 was replaced by TCF3-sgRNA-8XPP7. (E) Percentage of cells with genomic loci (IDR3 or TCF3) and HP1α condensate association in EpiGo-Control (dCas9) or EpiGo-KRAB (dCas9-KRAB) cell lines. Data are presented as means ± SD (n=3). (F) Colocalization of IDR3, TCF3 and HP1α in EpiGo-Control (dCas9) or EpiGo-KRAB (dCas9-KRAB) cells. A pair of IDR3 and TCF3 loci were dual-labelled by expression of IDR3-sgRNA-8XPP7, PCP-GFP, TCF3-sgRNA-8XMS2, MCP-SNAP, and dCas9 or dCas9-KRAB. U2OS-HP1α-HaloTag cells was used to examine the colocalization among IDR3 (green), TCF3 (blue) and HP1α (red). (G) Percentage of cells with IDR3 and TCF3 colocalization in EpiGo-Control (dCas9) and EpiGo-KRAB (dCas9-KRAB) cell lines. Data are presented as means ± SD (n=3).

To test how efficiently EpiGo-KRAB mediates genomic interaction and HP1α association, we chose a pair of loci IDR3 and TCF3 on the p-arm of human chromosome 19, which have been described previously ^15^. As shown in **Fig. 1B**, IDR3 located close to 5’-UTR of the THEG gene and TCF3 loci located at intron 4 of the TCF3 gene. RNA-seq data of U2OS cells showed that THEG was silenced, while the TCF3 was active. ChIP-seq of H3K9me3 showed that the 50 kb Region 1 containing IDR3 was highly H3K9 trimethylated, while the 50 kb Region 2 containing TCF3 showed very low H3K9me3. The genomic distance of Region 1 and Region 2 is about 1.2 Mb. As shown in **Fig. 1C**, IDR3 was detected to associate with HPα condensates in EpiGo-Control cells expressed dCas9, IDR3-sgRNA-8XPP7 and PCP-GFP, possibly due to high endogenous H3K9me3 levels of Region 1. Similar colocalization of IDR3 and HP1α was observed in EpiGo-KRAB cells (dCas9-KRAB, IDR3-sgRNA-8XPP7 and PCP-GFP). Nevertheless, the percentage of cells with IDR3 and HP⍰ association increased from 53% to 81% (**Fig. 1E**). As seen in **Fig. 1D** and **1E**, almost no colocalization of TCF3 and HPα was detected in EpiGo-Control cells (dCas9), while there were 51% of cells shown their colocalization in EpiGo-KRAB cells (dCas9-KRAB). Live cell tracking showed dramatical decrease of the dynamics when the TCF3 associated with HP1α in EpiGo-KRAB cells (**Fig. S1** and **Movie S1-S4**). These results suggest that EpiGo-KRAB can efficiently relocate genomic loci to HP1α condensates. To confirm whether EpiGo-KRAB could mediate genomic interactions, we utilized a tricolor system to label IDR3 (sgRNA-8XPP7 and PCP-GFP), TCF3 (sgRNA-8XMS2 and MCP-SNAP) and HP⍰ (HaloTag) simultaneously in single cells. As shown in **Fig. 1F**, colocalization of IDR3, not TCF3 with HP⍰ condensates were confirmed in EpiGo-Control cells (dCas9). However, colocalization of IDR3, TCF3 and HP⍰ was observed in EpiGo-KRAB cells (dCas9-KRAB). The genomic interactions between IDR3 and TCF3 increase from less than 2% to 43% (**Fig. 1G**).

Interestingly, IDR3 in EpiGo-Control and EpiGo-KRAB cells (**Fig. 1C, 1F**) or TCF3 in EpiGo-KRAB cells (**Fig. 1D, 1F**) was found to associate, but not extensively overlap with HP1α condensates. This observation intrigues us re-examine the spatial arrangements of endogenous H3K9me3 and HP1α condensates. As shown of the widefield microscopy imaging data in **Fig. S2A** and **S2B**, H3K9me3 and HP1α condensates well overlapped, which have been reported previously ^16^. However, 3D-SIM data (**Fig. S2C** and **S2D**) showed the endogenous H3K9me3 and HP1α condensates association but with limited overlapping, which is consistent with our findings of specific spatial arrangement between IDR3 or TCF3 loci and HP1α condensates in EpiGo-KRAB cells.

Long-range genomic interaction mediated by H3K9me3 occur at the hundreds of kilobases to megabase scales from Hi-C heatmap, which leads to hypothesizing the potential role of H3K9me3 in genome compartmentalization ^6^. Here we tried to visualize H3K9me3-mediated genome organization at the megabase scale in live cells. To induce H3K9me3 at the megabase scale, we mined the chromosome-specific repeats across hundreds of kilobases to megabase regions of human genome. We found a repeat class which consists of 836 copies of CRISPR target sites spanning ~17 megabases at the q-arm of chromosome 19, and we dubbed it C19Q as EpiGo targets (**Fig. 2A** and **Table S1**). C19Q can be visualized by co-expression of dCas9-KRAB, sgRNA-2XPP7 and PCP-GFP. We termed this suite of reagents as EpiGo-C19Q-KRAB. EpiGo-C19Q-KRAB could presumably recruit SETDB1 ^17^ to C19Q genomic sites which deposits loci-specific H3K9me3 ^13^. HP1α could then interact with loci or regions with H3K9me3 ^18^. Thus, we expect that EpiGo of C19Q will allow us to track the change of genome reorganization upon epigenetic modification at a megabase scale.

**Fig. 2.**
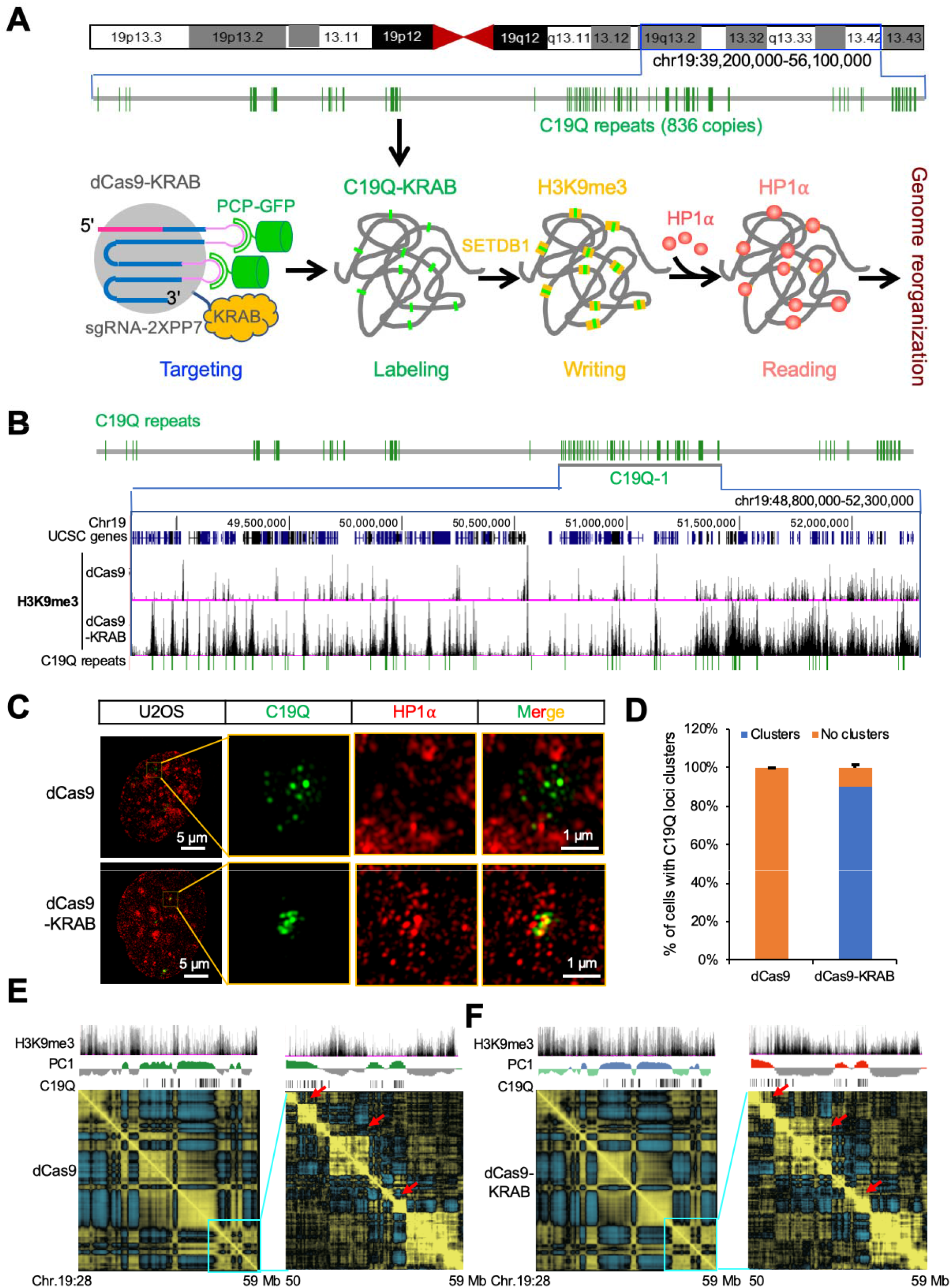
Live imaging of genome reorganization upon H3K9me3 by EpiGo-KRAB. (A) A scheme of the EpiGo-C19Q-KRAB system. EpiGo-C19Q-KRAB consists of dCas9-KRAB, PCP-GFP and sgRNA-2XPP7 targeting to C19Q repeats, which contain 836 copies of target sites on the q-arm of chromosome 19. EpiGo-C19Q-KRAB will recruit SETDB1 and induce H3K9me3 at each target sites. H3K9me3 regions will recruit HP1α and induce genome reorganization. (B) H3K9me3 state of EpiGo-C19Q-Control (dCas9) and EpiGo-C19Q-KRAB (dCas9-KRAB) cell lines. ChIP-seq of H3K9me3 was performed in these two cell lines. C19Q-1 region (chr19:48,800,000-52,300,000) is chosen to show the difference of H3K9me3 state between EpiGo-C19Q-Control and EpiGo-C19Q-KRAB cell lines. (C) 3D-SIM images of C19Q and HP1α in EpiGo-C19Q-Control and EpiGo-C19Q-KRAB cells. C19Q (green) was labelled by GFP and HP1α (red) was labelled by HaloTag. The colocalization between C19Q and HP1α is shown in merged images. (D) Percentage of C19Q foci clusters in EpiGo-C19Q-Control (dCas9) and EpiGo-C19Q-KRAB (dCas9-KRAB) cell lines. Data are presented as means ± SD (n=3). (E) Hi-C heatmap shown the compartmentalization of the C19Q region in EpiGo-C19Q-Control (dCas9) cells. H3K9me3 state, PC1 and C19Q repeats (target sites) shown at the top of the Hi-C heatmap (Chr19: 28,000,000-59,000,000). Hi-C heatmap of a local region (chr19: 50,000,000-59,000,000) is used to show the local compartmentalization. (F) Hi-C heatmap shown the compartmentalization of the C19Q region in EpiGo-C19Q-KRAB (dCas9) cells. All the data processing and display are the same as (E). Red arrows show the regions with increased neighboring interactions in the heatmap.

U2OS-HP1α-HaloTag cells were used to generate cell lines stably expressed C19Q-sgRNA-2XPP7, PCP-GFP and dCas9 or dCas9-KRAB, resulting in U2OS-EpiGo-C19Q-Control or U2OS-EpiGo-C19Q-KRAB cells for direct visualization of long-range interactions among target sites of C19Q upon ectopic H3K9me3 modification. ChIP-seq analysis confirmed that most target sites of C19Q successfully acquired ectopic H3K9me3 (**Fig. 2B and Fig. S3**). Structured illumination microscopy (3D-SIM) was used to acquire high resolution imaging of C19Q and HP1α. As shown in **Fig. 2C**, discrete foci of C19Q (green) were visible by CRISPR labeling in EpiGo-C19Q-Control (dCas9) cells, which barely colocalize with HP1α (red). After induction of H3K9me3 by EpiGo-KRAB, C19Q foci clustered and decorated on the surface of HP1α condensates in EpiGo-C19Q-KRAB (dCas9-KRAB) cell lines, which confirmed the association but limited overlapping between H3K9me3 regions and HP1α condensates. Quantitative analysis confirmed that C19Q foci were not clustered in EpiGo-C19Q-Control cells, while 90% of C19Q foci showed clustering in EpiGo-C19Q-KRAB cells (**Fig. 2D**). C19Q loci were highly dynamic in EpiGo-C19Q-Control cells but the mobility of C19Q loci dramatically decreased in EpiGo-C19Q-KRAB cell lines (**Fig. S4** and **Movie S5-S6**). Long-term live cell tracking of C19Q (green) confirmed that C19Q loci barely colocalized with HP1α (red) over time in EpiGo-C19Q-Control cells (dCas9) (**Fig. S5**). In EpiGo-C19Q-KRAB cell lines (dCas9-KRAB), However, C19Q loci dynamically interacted with HP1α and clustered at the surface of HP1α condensates, and eventually adjacent HP1α condensates coalesced together (**Fig. S5**). These results support a role of H3K9me3 regions as scaffolds in driving phase separation of HP1α droplets and large-scale genome organization.

To confirm long-range genomic interactions or clustering of C19Q loci in the EpiGo-KRAB system, Hi-C was performed on U2OS-EpiGo-C19Q-Control (dCas9) and U2OS-EpiGo-C19Q-KRAB (dCas9-KRAB) cell lines. In contrast to visualizing genomic interactions and clustering of C19Q loci from imaging in U2OS-EpiGo-C19Q-KRAB cell lines, there were little changes of overall interaction frequency between pairs of C19Q target sites in the above two cell lines (**Fig. S6**). No drastic changes of compartmentalization of the C19Q region were observed in the Hi-C matrix of the genome upon EpiGo-KRAB induction (**Fig. 2E** and **2F**), although there were substantial changes of H3K9me3 states. Focus on the local region (chromosome 19: 50-59 Mb), we have found a few regions with C19Q repeats (red arrows, **Fig. 2E** and **2F**) that show increased interactions with their neighbor regions, leading to compartment merging in U2OS-EpiGo-C19Q-KRAB (dCas9-KRAB) cells. Thus, Hi-C matrix detected EpiGo-KRAB induced local interactions but much less frequency of long-range interactions, while the microscopy methods observed more prominent long-range genomic interactions or clustering of C19Q loci in the EpiGo-KRAB system. It will be worthy to further investigate the differences of Hi-C and microscopy methods on the study of 3D genome in the diverse biological context.

Despite the long-observed correlation, whether epigenetic modifications can regulate 3D chromatin architecture remains poorly understood. The EpiGo system provides a powerful tool to allow epigenetic modifications at specific genes or regions and track their changes of location, structure and dynamics. Our data indicate that H3K9me3 mediated HP1α association and genomic clustering may disrupt existing chromatin compartments in a way that is analogous to the effects of CTCF and cohesin mediated loops. Drastic changes of epigenetic modifications such as H3K9me3, H3K27me3 and DNA methylation occur in stem cell differentiation, embryonic development and many diseases ^19–20^. With the EpiGo system, it will be intriguing to explore how tissue-specific epigenetic modifications are established and maintained at kilobase to megabase scales, and how they regulate chromatin architecture, gene expression and cell fate decision.

## Acknowledgements

We thank Luke Lavis (Janelia Research Campus, Howard Hughes Medical Institute, Ashburn, VA, USA) for the HaloTag JF-549. U2OS Genomic DNA was a gift Xingxu Huang. We thank Jianan Li for help with Lentivirus production. We are grateful to Guisheng Zhong, Cuiping Tian, Xiaoming Li and Yi Qian for help with imaging. We thank Pengwei Zhang and Shuangli Zhang for help with cell sorting. DeltaVision OMX™ V4 microscopy was provided by Shanghai Institute for Advanced Immunochemical Studies (SIAIS) at Shanghaitech University and Fluorescence activated cell sorting (FACS) was provided by iHuman Institute of ShanghaiTech. We thank members of the Ma and Xie’s laboratories for their comments during preparation of the manuscript. This work was funded by National Natural Science Foundation of China (No. 31970591 to H. Ma) and the Shanghai Pujiang program (19PJ1408000 to H. Ma). The initiation of this work in the Pederson laboratory was funded in part by U.S. National Institutes of Health grant U01 DA-040588 through the NIH 4D Nucleome Initiative.

## Author contributions

H.M. conceived this project. H.M. and Y.F. designed the experiments. A.N., and S.Z. performed data mining of chromosome-specific repeats. C.Y. construct plasmids. Y.F. performed fluorescence imaging and 3D-SIM. Y.W. performed RNA-seq, ChIP-seq and Hi-C. H.M., Y.F., W.X., Y.W., J.Z., X.X., and T.P. interpreted data. Y.F., Y.W., W.X. and H.M. wrote the paper with input from all the authors.

## Methods

### Cell culture

The U2OS (human bone osteosarcoma epithelial, female) cell, and HEK293T cells (human embryonic kidney epithelial, female) were cultured in DMEM (Life Technologies) with high glucose in 10% FBS (fetal bovine serum, Life Technologies). All cells were cultured at 37°C and 5% CO_2_ in a humidified incubator.

### Chromosome-specific repeats for the EpiGo system

Mining of chromosome-specific repeats was described previously with some modifications ^21^. The human reference genome (assembly GRC h37/hg19) was downloaded from the UCSC Genome Browser (http://genome.ucsc.edu) to find target regions and design guide RNAs. The bioinformatics tool Jellyfish ^22^ was used to find all 15-mers (12-mers ending with NGG or starting with CCN) with at least 5 copies in any chromosome. The 15-mers with more than 100,000 targets were filtered out. Each candidate 15-mer was searched for off-targets in all other chromosomes. The candidate 15-mer was discarded if there was any cluster of 5 off-targets or more within any 50 kb region. The C19Q repeats (**Table S1**), which consists of 836 copies of CRISPR target sites spanning ~17 megabases at the q-arm of chromosome 19, was chosen as EpiGo targets for the genome organization study. IDR3 and TCF3 target loci were described previously ^15^.

### Plasmids construction

The expression plasmid pHAGE-TO-dCas9 has been described previously ^14^, in which HSA-P2A was inserted at the N-terminal of dCas9 resulting in pHAGE-TO-HSA-P2A-dCas9 and KRAB was then subcloned into the C-terminal of dCas9 resulting pHAGE-TO-HSA-P2A-dCas9-KRAB. The expression plasmid pHAGE-EFS-PCP-GFP was described previously ^14^. pHAGE-EFS-MCP-SNAP was cloned by replacing GFP with SNAP. The expression vector for guide RNA was based on the pLKO.1 lentiviral expression system, in which TetR-BFP-P2A-sgRNA-2XPP7 or 8XPP7 was inserted right after the phosphoglycerate kinase (PGK) promoter, resulting pTetR-P2A-BFP-sgRNA-2XPP7 and pTetR-P2A-BFP-sgRNA-8XPP7. The expression plasmid for guide RNAs targeting to IDR3, TCF3 and C19Q was made using the rapid guide RNA construction protocol described previously ^21^. The dual-sgRNA expression vector pPUR-hU6-Sirius-8XPP7-mU6-Sirius-8XMS2 has been described previously ^15^. IDR3 and TCF3 were inserted into this vector, resulting pPUR-hU6-Sirius-IDR3-8XPP7-mU6-Sirius-TCF3-8XMS2.

The donor plasmid for knock-in of HaloTag at the C-terminal of HP1α was made by Golden Gate cloning. The donor consists of three fragments: A 400-bp fragment upstream of stop codon of HP1α (left arm), HaloTag coding sequences and a 400-bp fragment downstream of stop codon of HP1α (right arm). Left arm and right arm fragments were amplified from U2OS genomic DNA isolated with Cell DNA isolation mini kit (Vazyme). The HP1α-HaloTag donor was assembled into pDONOR vector by Golden Gate cloning method ^21^, resulting in pDONOR-HP1α-HaloTag. The guide RNA targeting sequence AAACAGCAAAGAGCTAAAGG spanning the stop codon (TAA, underlined) of HP1α was cloned into guide RNA expression vector pLH-sgRNA2 ^14^, resulting in pLH-sgRNA2-HP1α.

### Generation of U2OS-HP1α-HaloTag cell lines

For knock-in of HaloTag into HP1α locus, U2OS cells were co-transfected with 200 ng of pHAGE-TO-Cas9, 600 ng of pLH-sgRNA1-HP1α and 600 ng pDONOR-HP1α-HaloTag using Lipofectamine 2000 (Life Technologies) for 6 hours and then replaced culture media containing 2 nM HaloTag-JF549. The transfected cells were cultured for additional 24-48 hours before examining the knock-in efficiency by fluorescent imaging or flow cytometry. Fluorescence imaging was used to check the proper localization of HP1α-HaloTag. The HaloTag positive cells was sorted by BD FACS Aria III equipped with 561 nm excitation laser, and the emission signals were detected by using filter at 610/20 nm (wavelength/bandwidth) for HaloTag-JF549. The localization of HP1α-HaloTag was examined again under fluorescence microscope after cultured for additional two weeks. The resulting cell line was named U2OS-HP1α-HaloTag.

### Generation of U2OS-EpiGo cell lines

For generation of U2OS-EpiGo cell lines, lentivirus for PCP-GFP, sgRNA, dCas9, or dCas9-KRAB was used. HEK293T cells were seeded into 6-well and transfected with 0.5 μg pCMV-dR8.2-dvpr, 0.3 μg pCMV-VSV-G and 1.5 μg of transfer plasmids (pHAGE-EFS-PCP-GFP, pTetR-P2A-BFP-sgRNA-2XPP7-C19Q, pHAGE-TO-HSA-P2A-dCas9 or pHAGE-TO-HSA-P2A-dCas9-KRAB) using Lipofectamine 2000 (Invitrogen) following manufacturer’s protocol. After 48 hours, supernatant was harvested and filtered with 0.45 μm polyvinylidene fluoride Syringe filters. The filtered supernatant was titrated, and either directly used or concentrated. The concentration was performed by using Lentivirus Concentration Reagent (Biodragon-immunotech. Inc). The concentrated viral particles were immediately used for transduction or stored at −80°C in aliquots.

U2OS-HP1α-HaloTag cells were grown to 40% confluency in 6-well plates and viral supernatants for PCP-GFP, sgRNA targeting to C19Q, dCas9 or dCas9-KRAB were added for transduction. Fresh media containing 500 ng/ml doxycycline (Sigma-Aldrich) was used 48 hours post-infection. After additional 24 hours, Alexa Fluor^®^ 647 anti-mouse CD24 Antibody (BioLegend) for HSA were added before cell sorting. Expression levels of PCP-GFP, sgRNA (BFP) and dCas9 (AlexaFluor 647) or dCas9-KRAB (AlexaFluor 647) was examined by BD FACS Aria III equipped 405 nm, 488 nm, and 647 nm excitation lasers, and the emission signals were detected by using filters at 450/40 nm (wavelength/bandwidth) for BFP, 530/30 nm for GFP and 662/20 nm for AlexaFluor 647. To generate single colonies, single cells with different levels of PCP-GFP, sgRNA and dCas9 or dCas9-KRAB were sorted into 96-well plates. After 10-14 days, triple positive colonies were further examined for CRISPR-based labeling of C19Q foci under fluorescence microscope. The C19Q labeled cells were propagated and named U2OS-EpiGo-C19Q-Control (PCP-GFP, C19Q sgRNA and dCas9) and U2OS-EpiGo-C19Q-KRAB (PCP-GFP, C19Q sgRNA and dCas9-KRAB) respectively.

### Plasmid transfection

For transfection, typically 20 ng each of PCP-GFP and/or MCP-SNAP plasmids, 1 μg of guide RNA plasmids and 200 ng dCas9 or dCas9-KRAB were co-transfected using Lipofectamine 2000. 3 days after transfection, cells were incubated with 2 nM HaloTag-JF549 and 5 nM SNAP-Cell^®^ 647-SiR (NEB) before imaging.

### Live cell imaging

All live cell imaging was carried out on a DeltaVision OMX™ V4 imaging system (GE Healthcare), equipped with a 60x (NA 1.42) Plan Apo oil-immersion objective (Olympus). The cells were cultured on No. 1.0 glass bottom dishes (MatTek). The microscope stage incubation chamber was maintained at 37°C and 5% CO_2_. HaloTag-JF549 was excited at 561 nm, and its emission was collected using filter at 609/37 nm (wavelength/bandwidth), GFP was excited at 488 nm and collected using filter at 498/30 nm (wavelength/bandwidth). Imaging data were acquired by DeltaVision Elite imaging (GE Healthcare Inc.) software. Tracking C19Q cluster formation was performed for 4 hours and images were captured every 30 minutes. For the representative images, the raw data were deconvoluted by softWoRx (GE Healthcare Inc.) software.

### 3D-SIM procedure

3D-SIM was performed on a DeltaVision OMX™ V4 system (GE Healthcare) equipped with a 60x (1.42 NA) Plan Apo oil-immersion objective (Olympus) and six lasers (405, 445, 488, 514, 568 and 642nm). Image stacks were captured and were reconstructed using softWoRx (GE Healthcare). Images were registered with alignment parameters obtained from calibration measurements with 100 nm diameter TetraSpeck Microspheres with four colors (Molecular Probes).

### ChIP-seq

Cells were collected and crosslinked by 1% formaldehyde. Cells were suspended in cell lysis buffer B (50 mM Tris-HCl PH 8.0, 20 mM EDTA, 0.3% SDS, freshly added protease inhibitor) for 10 min on ice before sonication. 1 mg beads (Invitrogen) were washed with PBS+5 mg/mL BSA for three times and incubated with 5 μg antibody at 4°C for 6-8 hours before adding chromatin. Sonicated chromatin was diluted in dilution buffer (16.7 mM Tris-HCl PH 8.0, 1.1% Triton X-100, 1.2 mM EDTA, 167 mM NaCl, freshly added protease inhibitor), and was incubate with pre-treated beads and antibodies by rotation at 4°C overnight. Then beads were washed with washing buffer (50 mM Hepes PH 8.0, 1% NP-40, 0.7% DOC, 0.5 M LiCl, freshly added protease inhibitor) for 5 times followed by wash with TE once. 100μL elution buffer (50 mM Tris-HCl pH 8.0, 1 mM EDTA, 1% SDS) was added and beads were incubated in Thermo mixer at 65°C for 30 min with max speed. Supernatant was collected and treated with protease K at 55°C for 2 hours, then purified with DNA purification kit (Tiangen). RNA was removed by treatment of RNase at 37°C for 1 hour. DNA was then purified by AMPure Beads (Beckman) and subjected to DNA library preparation as described below.

### RNA-seq library preparation and sequencing

5 mg RNA was extracted using quick-RNA MiniPrep kit (Zymo) and then treated with DNase I (Fermentas) at 37°C for 1 hour. RNA was then purified using AMPure beads. Poly-A tailed mRNA was collected using Dynabeads™ mRNA purification kit (Invitrogen). Purified RNA was fragmented with RNA Fragmentation Buffer (NEB) at 95°C for 5 min. Reaction was stopped and RNA was purified by AMPure beads. First strand cDNA was synthesized with a commercial kit using both oligo dT and random primers (Invitrogen). Second strand cDNA was synthesized with second strand synthesis buffer (Invitrogen), MgCl_2_, DTT, dNTP, dUTP, RNase H (Fermentas), E. coli DNA ligase (NEB) and DNA polymerase I (NEB). DNA was purified after 2 hours incubation on thermomixer at 16°C. Synthesized cDNA was subjected to DNA library preparation as described below.

### Library construction

Purified DNA or cDNA was subjected to NEBNext Ultra II DNA Library Prep Kit for Illumina (NEB). The DNA was resuspended in 25 μl ddH2O, followed by end-repair/A-tailing with 3.5 μl End Prep buffer and 1.5 μl enzyme mix according to manufactory instruction. The ligation reaction was then performed by adding diluted 1.25 μl adaptors (NEB), 15 μl Ligation master mix, 0.5 μl Ligation enhancer, at 4°C for overnight. The ligation reaction was treated with 1.5μl USERTM enzyme according to the instruction and was purified by AMPure beads. The 1^st^ round PCR was performed by adding 25 μl 2x KAPA HiFi HotStart Ready Mix (KAPA biosystems) with primers of NEB Oligos kit, with the program of 98°C for 45 s, (98°C for 15 s and 60°C for 10 s) with 8-9 cycles and 72°C for 1 min. DNA was purified using 1x AMPure beads and a 2^nd^ round PCR was performed with 25 μl 2x KAPA HiFi HotStart Ready Mix with Illumina universal primers, and same PCR program for 8 cycles. The final libraries were purified by 1x AMPure beads and subjected to next-generation sequencing. All libraries were sequenced on Illumina HiSeq 2500 or HiSeq XTen according to the manufacturer’s instruction.

### sisHi-C library generation and sequencing

The sisHi-C library generation was performed as described previously ^23^. Briefly, spermatogenetic cells were fixed with 1% formaldehyde at room temperature (RT) for 10 min. Formaldehyde was quenched with glycine for 10 min at RT. Cells were washed with 1XPBS for two times and then lysed in 50 μl lysis buffer (10 mM Tris-HCl pH7.4, 10 mM NaCl, 0.1 mM EDTA, 0.5% NP-40 and proteinase inhibitor) on ice for 50 min. After spinning at 3000 rpm/min in 4°C for 5 min, the supernatant was discarded with a pipette carefully. Chromatin was solubilized in 0.5% SDS and incubated at 62°C for 10 min. SDS was quenched by 10% Triton X-100 at 37°C for 30 min. Then the nuclei were digested with 50 U Mbo I at 37°C overnight with rotation. Mbo I was then inactivated at 62°C for 20 minutes. To fill in the biotin to the DNA, dATP, dGTP, dTTP, biotin-14-dCTP and Klenow were added to the solution and the reaction was carried out at 37°C for 1.5 hours with rotation. The fragments were ligated at RT for 6 hours with rotation. This was followed by reversal of crosslink and DNA purification. DNA was sheared to 300-500 bp with Covaris M220. The biotin-labeled DNA was then pulled down with 10ul Dynabeads MyOne Streptavidin C1 (Life Technology). Sequencing library preparation was performed on beads, including end-repair, dATP tailing and adaptor-ligation. DNA was eluted twice by adding 20ul water to the tube and incubation at 66°C for 20 minutes. 9-15 cycles of PCR amplification were performed with Extaq (Takara). Finally, size selection was done with AMPure XP beads and fragments ranging from 200 bp to 1000 bp were selected. All the libraries were sequenced on Illumina HiSeq2500 or HiSeq Xten according to the manufacturer’s instruction.

### Imaging data analysis

The maximum projection of Z stack was used for the quantification of percentage of cell numbers (200 cells in total for each sample) with genomic loci interactions or clustering. Linescan analysis for the colocalization of HP1α and H3K9me3 was performed with Fiji software (Image J 1.52p). Trajectory of loci dynamics was analyzed by MTrack2 Plugins. Fluorescent puncta were identified as local maxima satisfying the minimum intensity and min/max peak with threshold by Gaussian fitting. Data are represented as mean ± SD. The exact number, n, of data points and the representation of n (cells, independent experiments) are indicated in the respective figure legends and in the Results.

### RNA-seq data processing

RNA-seq data were mapped to hg19 reference genome by Tophat. The gene expression levels were calculated by Cufflinks (version 2.2.1) using the refFlat database from the UCSC genome browser.

### ChIP data processing

The paired-end ChIP reads were aligned with the parameters: -t -q -N 1–L 25 -X 1000 --no-mixed --no-discordant. All unmapped reads, non-uniquely mapped reads, reads with low mapping quality (MAPQ < 20) and PCR duplicates were removed. For downstream analysis, we normalized the read counts by computing the numbers of reads per kilobase of bin per million of reads sequenced (RPKM) for 100-bp bins of the genome. To minimize the batch and cell type variation, RPKM values across whole genome were further Z-score normalized. To visualize the ChIP signals in the UCSC genome browser, we generated the RPKM values on a 100 bp-window base. H3K9me3 peaks were called using MACS2 ^24^ with the parameters -g hs -broad -nomodel -nolambda --broad and noisy peaks with very weak signals (RPKM < 5) were removed from further analyses. Adjacent peaks with very close distance (<5kb) were merged for downstream analyses.

### Hi-C data mapping

Paired end raw reads of Hi-C libraries were aligned, processed and iteratively corrected using HiC-Pro (version 2.7.1b) as described ^25^. Briefly, sequencing reads were first independently aligned to the human reference genome (hg19) using the bowtie2 end-to-end algorithm and “-very-sensitive” option. To rescue the chimeric fragments spanning the ligation junction, the ligation site was detected and the 5’ fraction of the reads was aligned back to the reference genome. Unmapped reads, multiple mapped reads and singletons were then discarded. Pairs of aligned reads were then assigned to Mbo I restriction fragments. Read pairs from uncut DNA, self-circle ligation and PCR artifacts were filtered out and the valid read pairs involving two different restriction fragments were used to build the contact matrix. Valid read pairs were then binned at a specific resolution by dividing the genome into bins of equal size. We chose 100-kb bin size for examination of global interaction patterns of the whole chromosome, and 40-kb bin size to show local interactions and to perform TAD calling. Then the binned interaction matrices were normalized using the iterative correction method ^25,26^ to correct the biases such as GC content, mappability and effective fragment length in Hi-C data.

### Identification of conventional chromatin compartments and refined-A/B

Conventional chromatin compartments A and B were identified with a method described previously ^27^ with some modifications. The normalized 100 kb interaction matrices for each stage were used in this analysis. Firstly, the bins that have no interactions with any other bins were removed before expected interaction matrices were calculated. Observed/Expected matrices were generated using a sliding window approach ^28^ with the bin size of 400 kb and the step size of 100 kb. Principal component analysis was performed on the correlation matrices generated from the observed/expected matrices. The first principal component of the correlation matrices together with gene density were used to identify A/B compartments. In this analysis, the correlation matrices were calculated according to the interaction matrices separated according to the location of predicted centromeres for each chromatin, as the principal component often reflects the partitioning of chromosome arms ^27^. As for Refined-A/B compartment, the calling method were similar to the conventional chromatin compartments, with the matrices restricted to 10 Mb one-by-one instead of whole chromosome arms. The PC1 value generated by those restricted matrices were taken as local PC1. Juicebox was used to generate all chromosome-wide as well as the zoom-in views of interaction frequency heatmap in this study. Both the conventional compartment correlation heatmap and the Refined-A/B compartment heatmap were generated with Java TreeView according to the corresponding correlation matrices.

### Data availability

All data are available in the manuscript or the supplementary materials. Raw sequence reads of U2OS-EpiGo-C19Q-Control and U2OS-EpiGo-C19Q-KRAB experiments are deposited in NCBI sequence read archive (SRA) database. Plasmids are available on Addgene.

### Code availability

All custom code used in this study is available upon request.

## Supplementary Figures

**Fig. S1.**
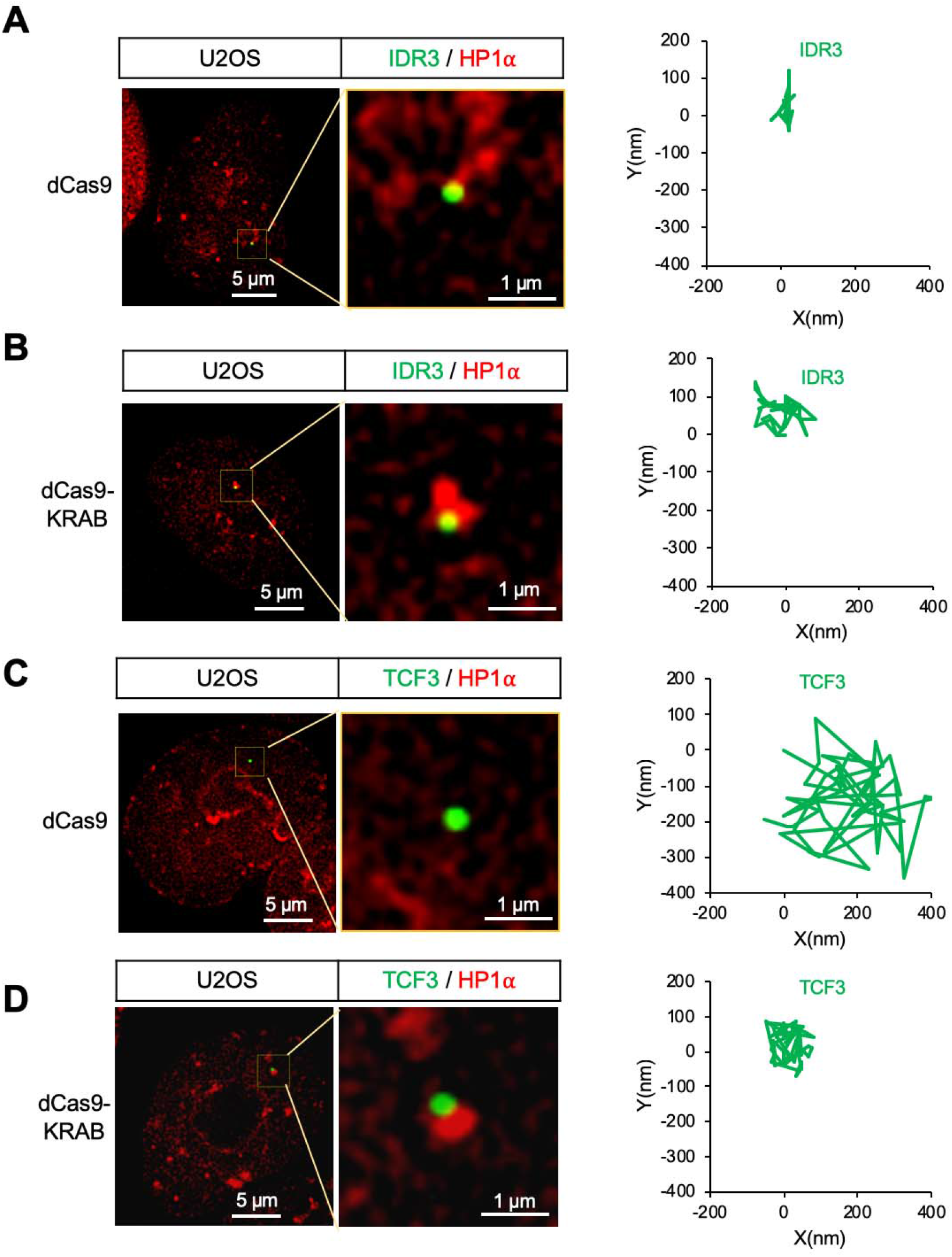
Tracking the Dynamics of IDR3 or TCF3 loci in EpiGo-KRAB cells. (A) Live cell tracking of IDR3 loci movement in EpiGo-Control cell lines. The dynamics of IDR3 (green) and HP1α (red) were tracked at 100 milliseconds per frame for 6 seconds (60 frames). All trajectories were shifted to start from the origin (0, 0) for easy comparison of the movement vectors. The locus movements were corrected for the possible movements of microscope stage. (B) Live cell tracking of IDR3 loci movement in EpiGo-KRAB cell lines. The image processing details are the same as described in (A). (C) Live cell tracking of TCF3 loci movement in EpiGo-Control cell lines. The image processing details are the same as described in (A) (D) Live cell tracking of TCF3 loci movement in EpiGo-KRAB cell lines. The image processing details are the same as described in (A).

**Fig. S2.**
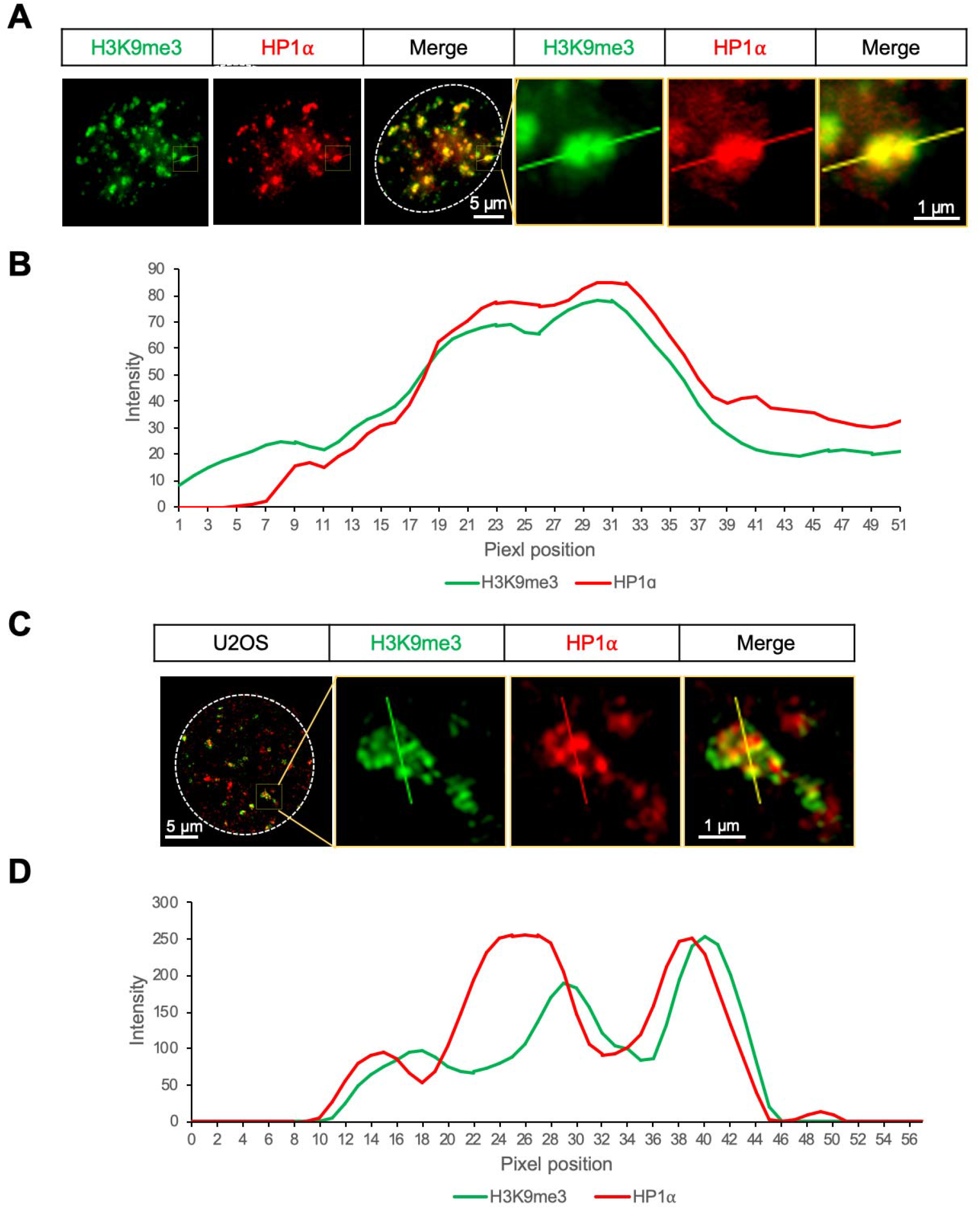
Colocalization of H3K9me3 and HP1α in U2OS cells. (A) Double staining of endogenous H3K9me3 and HP1α from widefield microscopy. H3K9me3 puncta (green) were detected by H3K9me3 antibody and HP1α (red) was tagged by HaloTag. (B) Linescan of the vertical line. linescan shows, from top to bottom, the intensities (arbitrary units) of H3K9me3 (green) and HP1α (red) along the line indicated in (A). (C) 3D-SIM images of endogenous H3K9me3 and HP1α. H3K9me3 puncta (green) were detected by H3K9me3 antibody and HP1α (red) was tagged by HaloTag. (D) Linescan of the vertical line. linescan shows, from top to bottom, the intensities (arbitrary units) of H3K9me3 (green) and HP1α (red) along the line indicated in (C).

**Fig. S3.**
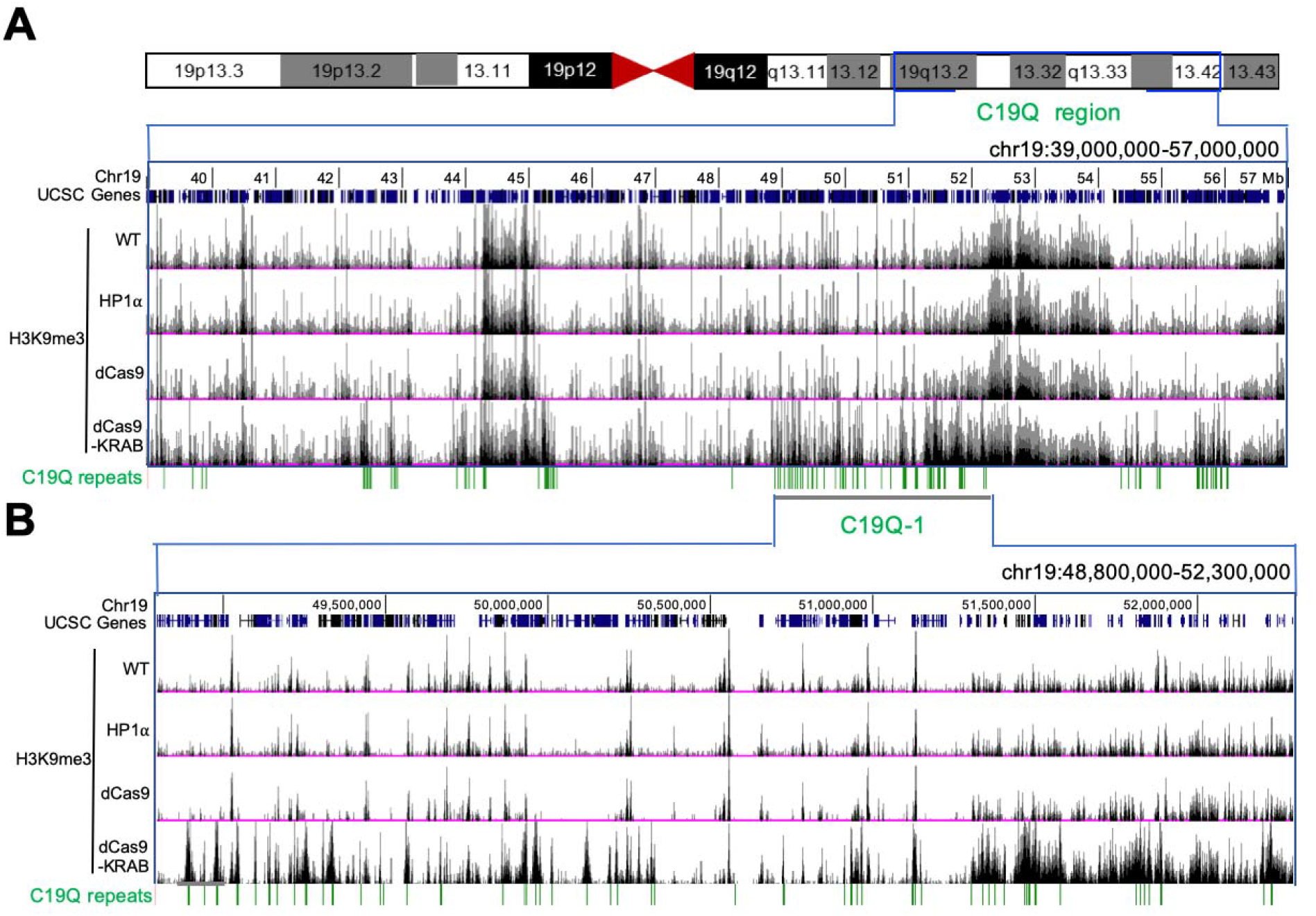
H3K9me3 of the C19Q region mediated by the EpiGo-KRAB system. (A) H3K9me3 states of the C19Q region in the cell lines of U2OS wildtype (WT), U2OS-HP1α HaloTag (HP1α), EpiGo-Control (dCas9) or EpiGo-KRAB (dCas9-KRAB). ChIP-seq of H3K9me3 was performed in these four cell lines. C19Q repeats shown the C19Q target sites. (B) C19Q-1 region is chosen to show the difference of H3K9me3 state between U2OS wildtype (WT), U2OS-HP1α-HaloTag (HP1α), EpiGo-C19Q-Control (dCas9) or EpiGo-C19Q-KRAB (dCas9-KRAB) cell lines. C19Q repeats shown the C19Q target sites.

**Fig. S4.**
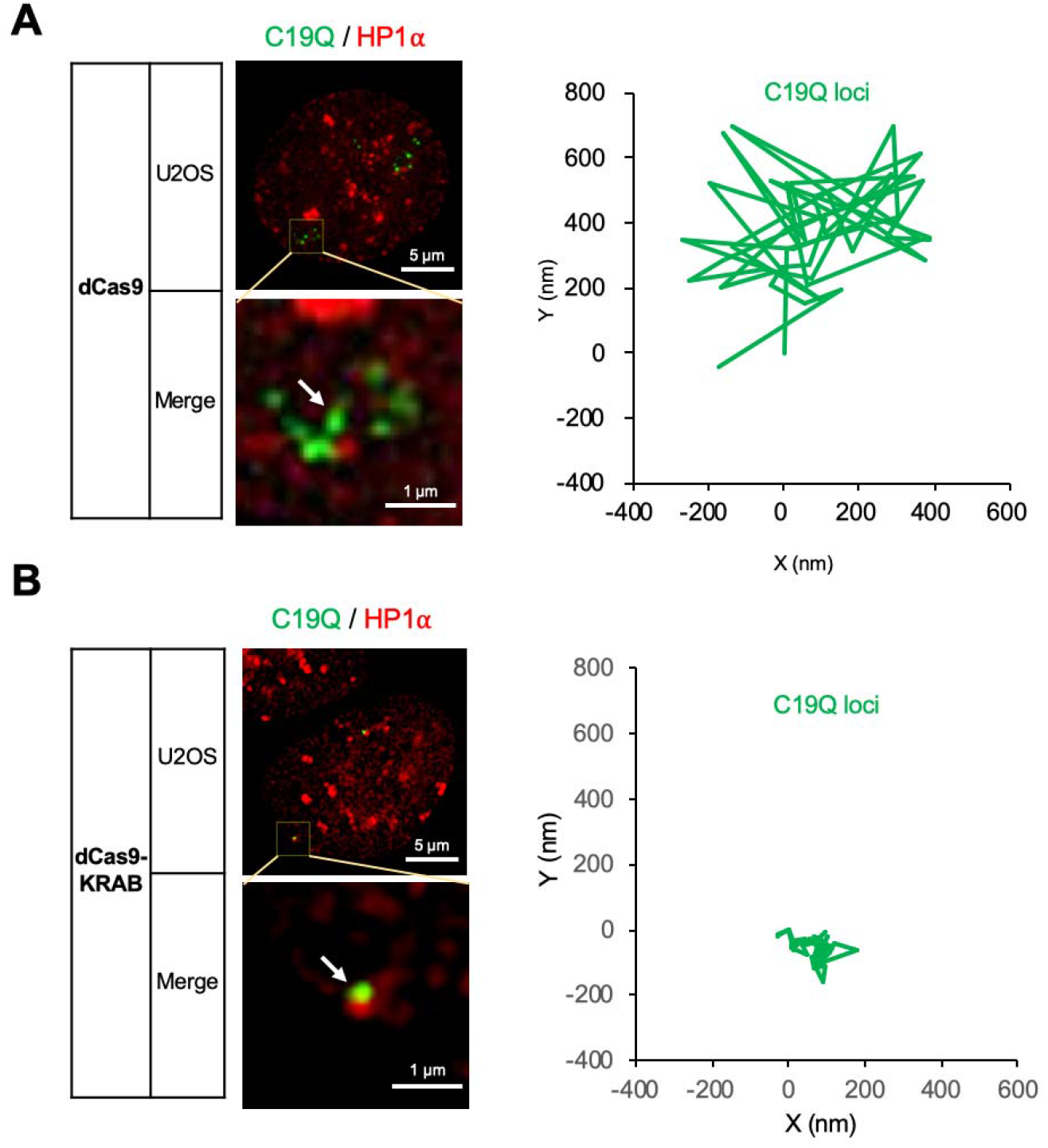
Tracking the Dynamics of C19Q loci in EpiGo-KRAB cells. (A) Live cell tracking of C19Q loci movement in EpiGo-C19Q-Control cell lines. The dynamics of C19Q (green) and HP1α (red) were tracked at 100 milliseconds per frame for 6 seconds (60 frames). The movement of locus C19Q loci (arrowed) were recorded. All trajectories were shifted to start from the origin (0, 0) for easy comparison of the movement vectors. The locus movements were corrected for the possible movements of microscope stage. (B) Live cell tracking of C19Q loci movement in EpiGo-C19Q-KRAB cell lines. The image processing details are the same as described in (A).

**Fig. S5.**
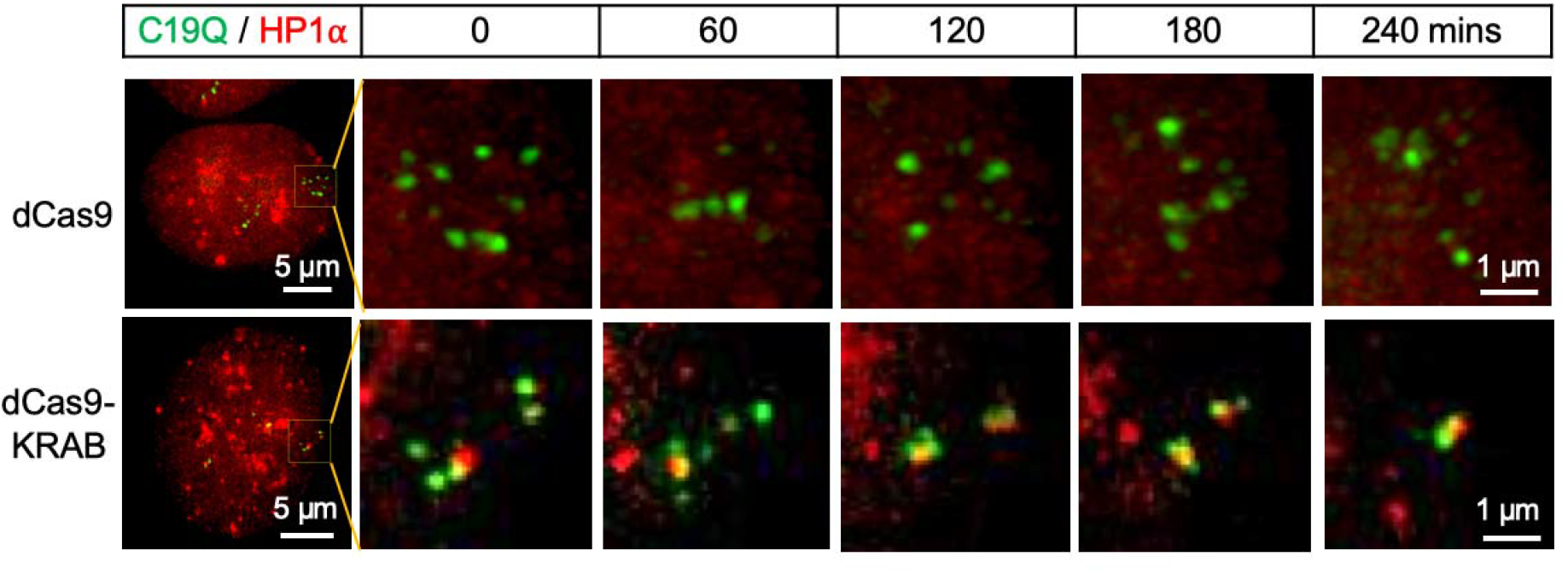
Live cell tracking of C19Q loci clustering in U2OS-EpiGo cell lines. EpiGo-C19Q-Control (dCas9) and EpiGo-C19Q-KRAB (dCas9-KRAB) cell lines were used for live cell tracking of C19Q loci clustering. The colocalization between C19Q (green) and HP1α (red) was shown at different time points indicated. The dCas9 or dCas9-KRAB was induced to express 24 hours before tracking. The dynamics of C19Q (green) and HP1α (red) were tracked for 240 mins. Images were captured every 30 mins.

**Fig. S6.**
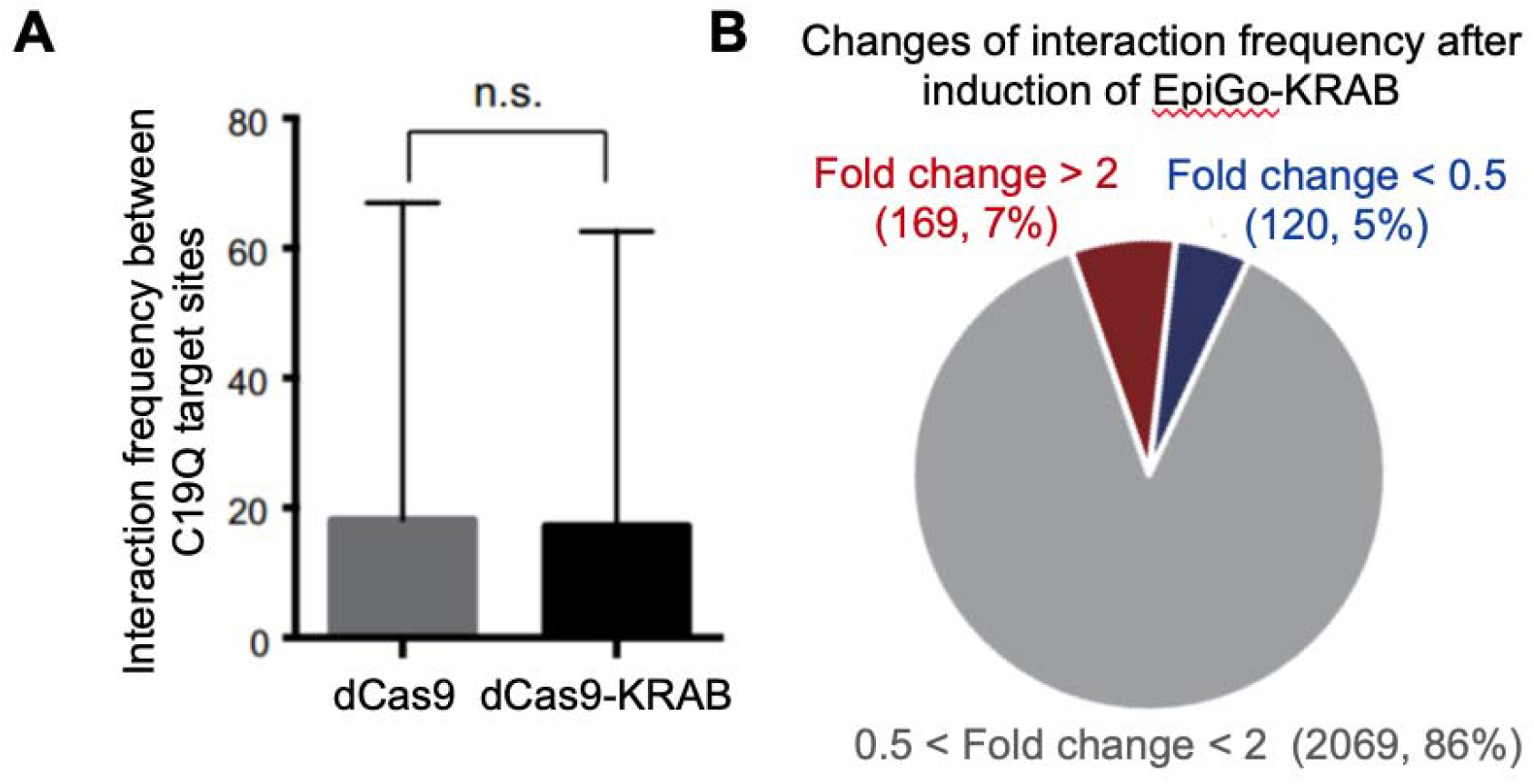
Interaction frequency between C19Q target sites shown in the Hi-C. (A) Interaction frequency between C19Q target sites in Hi-C matrix from EpiGo-C19Q-Control (dCas9) and EpiGo-C19Q-KRAB (dCas9-KRAB) cell lines were compared. n.s.: no significance. (B) Changes of interaction frequency after induction of EpiGo-KRAB in EpiGo-C19Q-KRAB (dCas9-KRAB) cell lines. The numbers of interactions in Hi-C matrix between EpiGo-C19Q-Control (dCas9) and EpiGo-C19Q-KRAB (dCas9-KRAB) cell lines were compared. The number of interactions increased more than 2 folds (>2) or decreased more than 2 folds (<0.5) was shown in brown or dark blue in pie chart. The Change of interaction numbers within 0.5 to 2 folds was shown grey in pie chart.

## Supplementary Tables

**Table S1. Target sites of C19Q on the q-arm of human chromosome 19.**

## Supplementary Movies

**Movie S1. Tracking the movement of IDR3 loci in U2OS-EpiGo-Control cells.**

Images were cropped to 300X300 pixels and each video includes 200 frames (a total time of 20 seconds). The imaging rate is 100 milliseconds per frame and the play rate is 30 frames per second. The individual locus movements were corrected for the possible movements of microscope stage.

**Movie S2. Tracking the movement of IDR3 loci in U2OS-EpiGo-KRAB cells.**

Images were cropped to 275X275 pixels and each video includes 200 frames (a total time of 20 seconds). The image processing details are the same as described in the Video S1.

**Movie S3. Tracking the movement of TCF3 loci in U2OS-EpiGo-Control cells.**

Images were cropped to 300X300 pixels and each video includes 200 frames (a total time of 20 seconds). The image processing details are the same as described in the Video S1.

**Movie S4. Tracking the movement of TCF3 loci in U2OS-EpiGo-KRAB cells.**

Images were cropped to 275X275 pixels and each video includes 200 frames (a total time of 20 seconds). The image processing details are the same as described in the Video S1.

**Movie S5. Tracking the movement of C19Q loci in U2OS-EpiGo-C19Q-Control cells.**

Images were cropped to 275X275 pixels and each video includes 200 frames (a total time of 20 seconds). The image processing details are the same as described in the Video S1.

**Movie S6. Tracking the movement of C19Q loci in U2OS-EpiGo-C19Q-KRAB cells.**

Images were cropped to 275X275 pixels and each video includes 200 frames (a total time of 20 seconds). The image processing details are the same as described in the Video S1.

